# Tumor-Derived SPP1 Drives Immunosuppressive Macrophage Reprogramming in Gastric Peritoneal Carcinomatosis

**DOI:** 10.64898/2026.06.21.733605

**Authors:** Lilia Turcios, Neelima Hosamani, Ellen J. Beswick, Eric Ubil, Maria Carey, Joshua Leinwand, Sachiyo Nomura, Jun Yan, Mark B. Evers, Joseph Kim, Mautin Barry-Hundeyin

## Abstract

Peritoneal carcinomatosis is a major cause of death in gastric cancer, yet effective therapies remain limited. Tumor-derived soluble factors are increasingly recognized as key regulators of the peritumoral microenvironment. Here, we nominate osteopontin (SPP1) as a tumor-derived mediator that orchestrates macrophage-driven immunoregulation in gastric peritoneal carcinomatosis. Using integrated analyses of human clinical datasets and murine models, we demonstrate that tumor-secreted SPP1 promotes macrophage recruitment and induces tolerogenic IL-10 production. Clinically, SPP1 correlated with inferior overall survival and progression-free survival in gastric cancer. In syngeneic murine models of gastric peritoneal carcinomatosis, intracavitary pharmacologic inhibition of SPP1 restricted peritoneal dissemination, impaired macrophage infiltration and suppressed IL-10 production. Consistent with these findings, macrophage depletion phenocopied antitumor effects of SPP1 inhibition, resulting in decreased metastatic burden. Collectively, these findings define a mechanism of tumor-macrophage crosstalk that promotes peritoneal dissemination and provide a rationale for therapeutic targeting of SPP1 in gastric peritoneal carcinomatosis.

## Introduction

Gastric peritoneal carcinomatosis represents a devastating manifestation of advanced gastric cancer and is associated with 2% long term survival rate and median overall survival less than one year^1,2^. This late-stage, therapy-refractory disease commonly manifests as metastatic peritoneal tumor implants, malignant ascites, or both. It is characterized by a profoundly immunosuppressive tumor microenvironment, stromal remodeling, and impaired blood-peritoneal barrier limiting drug delivery^3,4^. Despite advances in chemoimmunotherapy, effective treatments for peritoneal carcinomatosis remain limited. Therefore, a deeper understanding of the cellular and molecular mechanisms that drive metastatic progression within the peritoneal tumor microenvironment (TME) is urgently needed.

Tumor-derived soluble mediators are increasingly recognized as key regulators of immune cell behavior within the TME^5–8^. Our prior work has shown that macrophage behavior is shaped by diverse inflammatory signals that sustain pro-tumorigenic networks^9,10^. Although intraperitoneal macrophages are abundant in gastric peritoneal carcinomatosis^11–13^, the tumor-derived factors that drive macrophage recruitment and functional reprogramming in have not been fully elucidated.

Osteopontin (transcript encoded by *SPP1*, protein referred to as OPN) is an integrin binding protein secreted by diverse cell types, associated with pro-tumorigenic macrophage states^14–20^ and autocrine control of epithelial mesenchymal transition^21^. Although it has been reported in observational studies as an independent predictor of gastric peritoneal relapse,^22^ its role in regulating tumor-macrophage interactions within the peritoneal metastatic niche remains undefined. Here, we demonstrate that gastric cancer soluble mediators promote anti-inflammatory macrophage polarization. SPP1 was highly expressed and secreted by gastric tumor cells, effectively recruiting macrophages to the TME and promoting tumor-promoting IL-10 phenotype. Clinically, *SPP1* correlated with worse overall survival and progression free survival. Targeting SPP1 in murine models of gastric peritoneal metastasis restricted peritoneal carcinomatosis. In addition, macrophage depletion phenocopied SPP1 inhibition by limiting peritoneal dissemination. Therefore, we identify tumor derived SPP1 is a key regulator of macrophage recruitment and immunosuppressive programming, revealing a tumor-macrophage signaling axis that drives metastatic progression in gastric peritoneal carcinomatosis.

## Results

### Gastric tumor-derived soluble mediators promote anti-inflammatory macrophage reprogramming

Our prior studies have shown that macrophage behavior is impacted by diverse inflammatory signals that maintain pro-tumorigenic networks^9,10^. Therefore, we postulated that tumor cells exploit soluble mediators to reprogram macrophages and establish a microenvironment that supports oncogenesis. To test this hypothesis, we examined the effects of murine YTN16 gastric secreted factors on macrophage phenotype. We performed integrated transcriptomic and protein-level analysis of bone marrow derived macrophages (BMDMs) cultured in YTN16 tumor cell-conditioned media (CM) compared with untreated BMDMs. We observed downregulation of genes associated with proinflammatory macrophage phenotype (*Il1b, Trem1*, *Cxcl1*, *Ccl3*) and antigen presentation (*Cd1d1*, *Cd33*, *Cd74*). In contrast, expression of genes associated with an anti-inflammatory phenotype (*Trem2, Mrc1, Pparg*) and function to dampen adaptive responses (*Arg1*, *Pd-l2*, *Cd200r1*) were upregulated with exposure to CM (Figure 1A). PPAR-RXR signaling pathway, a critical promoter of M2-like polarization, was upregulated with CM exposure (Figure 1B). Gene ontology enrichment analysis identified significant upregulation of IL-10 associated pathways with macrophage CM treatment (Figure 1C).

**Figure 1:**
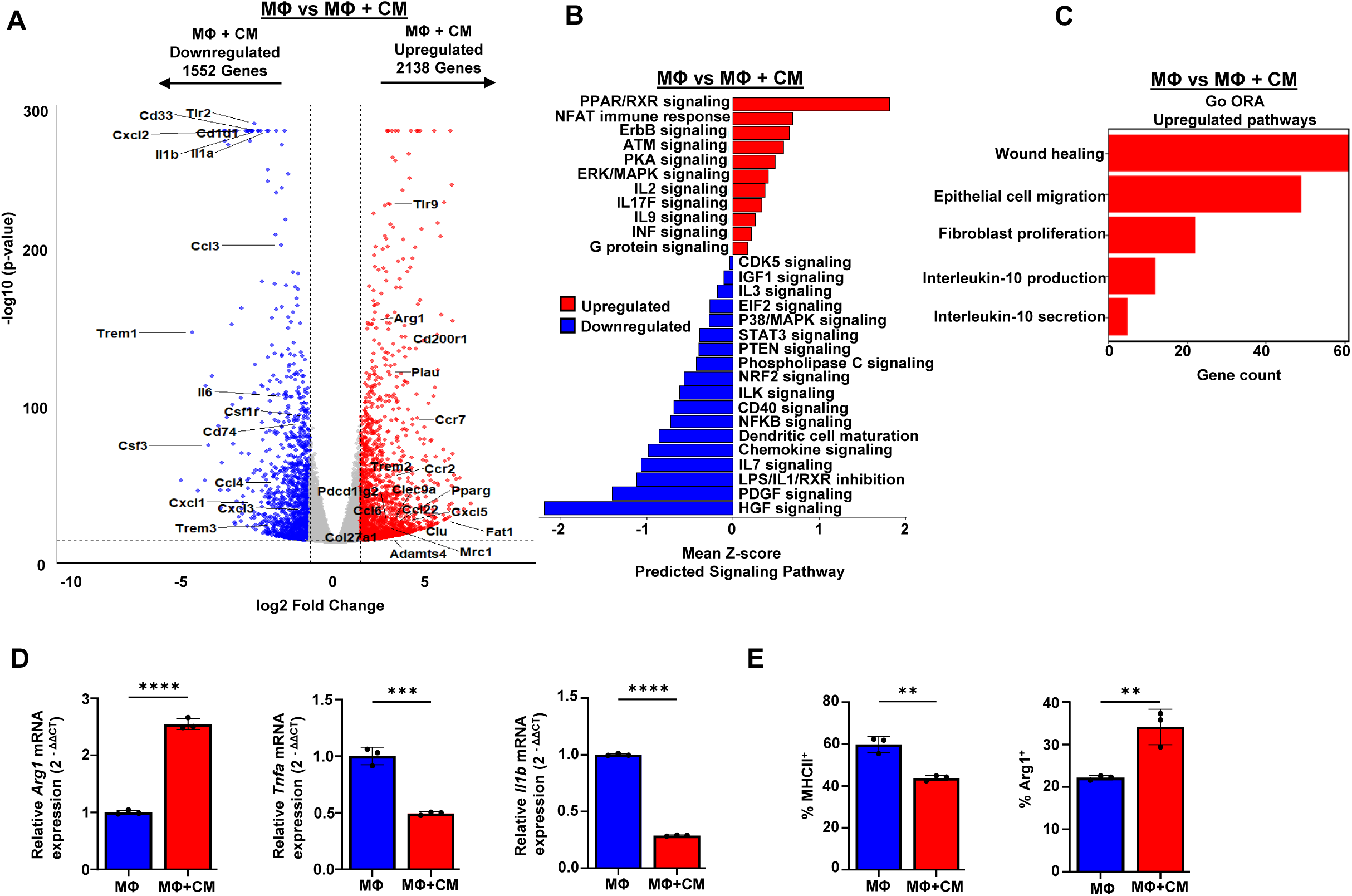
Gastric cancer secreted factors promote an anti-inflammatory macrophage phenotype. (A-C) Differential gene expression analysis (A), predicted signaling pathways (B) and GSEA (C) of bone marrow derived macrophages (BMDMs, MՓ) treated with tumor cell-conditioned media (CM) compared with untreated BMDMs. (D) RT-qPCR analysis of *Arg1*, *Tnfa*, *Il1b* expression in BMDMs treated with CM relative to control BMDMs. (E) Flow cytometry showing protein levels of MHCII and Arg-1 in BMDMs treated with CM compared with untreated BMDMs. Both treated and untreated BMDMs were stimulated with LPS. Data are representative of N=3 per group. \*\**p< 0.01*; \*\*\**p < 0.001*, \*\*\*\**p< 0.0001*, by unpaired t test was used to compare the differences between groups.

We validated our observations by performing RT-qPCR to measure shifts in transcript abundance and by using flow cytometry and cytokine analysis to evaluate whether these transcript-level changes were reflected at the protein level. BMDM exposure to CM induced transcriptional activation of *Arg1* and attenuated *Tnfα* and *Il1β* expression compared to untreated BMDM (Figure 1D). Accordingly, BMDM treatment with CM resulted in diminished expression of MHCII and increased expression of Arg-1 by flow cytometry (Figure 1E). Findings were recapitulated using RAW 267.4 macrophages. Treatment of BMDM with CM resulted in a decrease gene expression of *Tnfα* and *Nos2* and an increase in *Arg1* compared to untreated macrophages. (Supplemental Figure1A). Flow cytometric analysis of RAW 267.4 macrophages revealed a decrease TNFα, MHCII, and CD86 expression and an increase in Arg-1 expression (Supplemental Figure 1B). Thus, we determined that soluble mediators released by gastric neoplastic cells promote anti-inflammatory macrophage reprogramming, including tolerogenic IL-10 production and secretory pathways.

### SPP1 is a clinically relevant candidate gastric tumor-secreted factor

To identify clinically relevant secreted factors with immunomodulatory potential, we analyzed differentially expressed genes in gastric cancer for those encoding secreted proteins in the UALCAN, GTex, TCGA, GCR, GSE66229, and GSE162214 databases. Notably, we identified *SPP1* among the top 5 differentially expressed genes in gastric cancer (Figure 2A). SPP1 was overexpressed in primary gastric cancer compared to normal gastric tissue in Western (TCGA, GCR) and Asian (ACRG) cohorts, indicating importance across diverse populations (Figure 2B-C). Similar transcript levels of *SPP1* were detected in both malignant ascites and primary tumors, suggesting persistent expression during metastasis (Figure 2D). In line with these findings in patient samples, we detected SPP1 at the transcript and protein levels in the murine model of peritoneal carcinomatosis (Figure 2E). SPP1-encoding transcripts were found in both peritoneal tumor tissue and malignant ascites by bulk RNA sequencing (Figure 2F), and western blot analysis confirmed robust SPP1 protein expression in peritoneal metastatic tumor tissue and cultured YTN16 cells (Figure 2G). Finally, SPP1 was abundant in CM from YTN16 cells, indicating secretion by neoplastic cells (Figure 2H).

**Figure 2:**
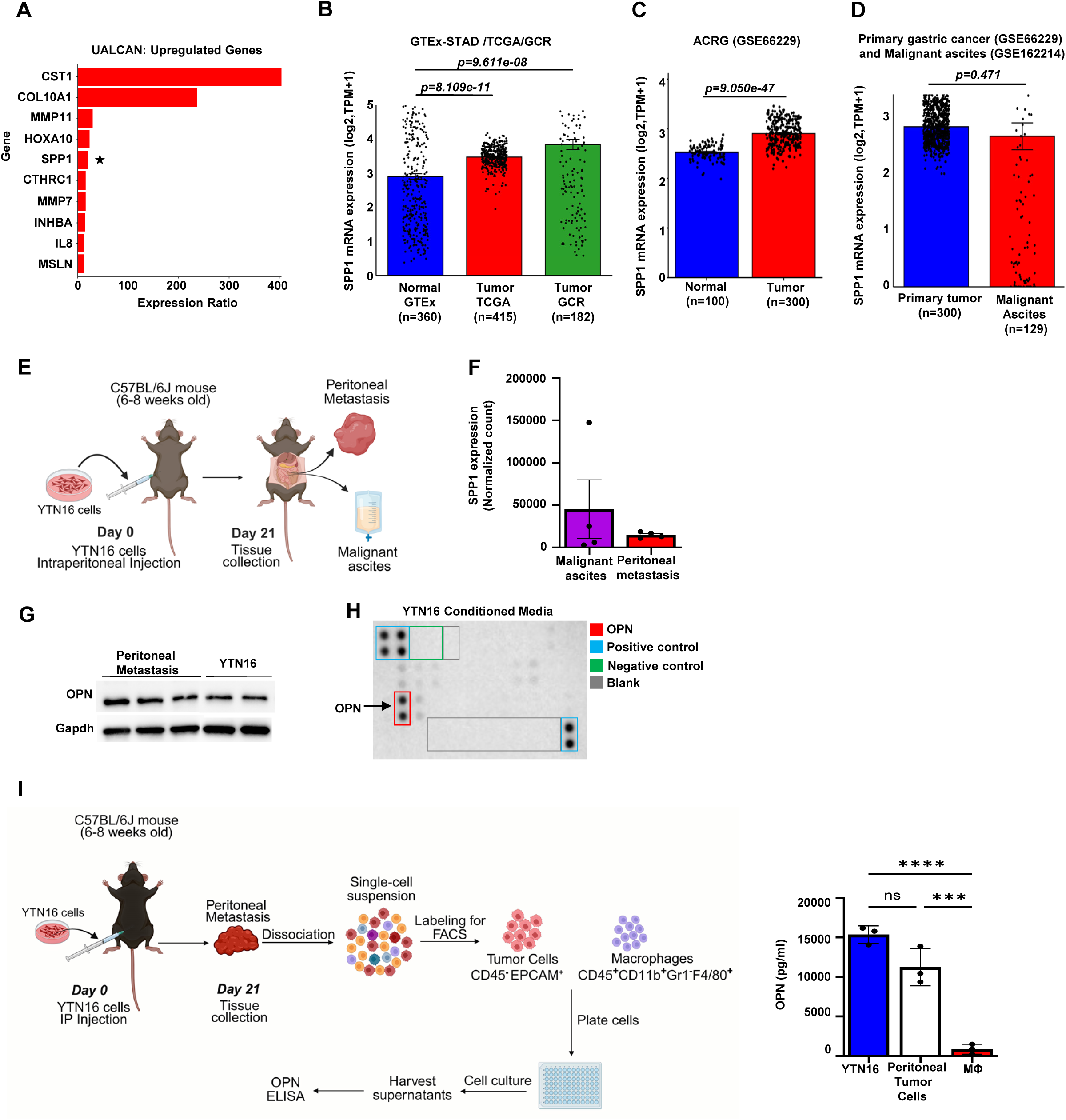
Gastric tumor cells are major secretors of SPP1. (A) Top 10 upregulated genes in gastric cancer adapted from the UALCAN database. (B) *SPP1* mRNA expression in gastric cancer compared with normal gastric tissue using TCGA-STAD and Gastric Cancer registry (GCR) datasets relative to normal gastric epithelium from the GTEx database (C) Comparison of *SPP1* gene expression in gastric cancer and normal gastric epithelium from the ACRG database (D) Comparison of SPP1 gene expression between primary gastric cancer and malignant ascites. (E) Schematic of murine gastric peritoneal carcinomatosis model created by intraperitoneal (IP) injection of syngeneicYTN16 gastric cancer cells. (F) Bulk RNA sequencing of SPP1 expression in murine peritoneal metastatic nodules and malignant ascites. (G) Western blot analysis of SPP1 protein expression in murine gastric peritoneal metastatic nodules and YTN16 murine gastric cell lines. (H) Multiplex cytokine assay from conditioned media from YTN16 murine gastric cancer cell line demonstrating SPP1 (OPN) secretion. (I) Schematic of ex-vivo sorting and culture of tumor cells (CD45^-^Epcam^+^) and macrophages (MՓ, CD45^+^CD11b^+^Gr1^-^F4/80^+^) from murine gastric peritoneal carcinomatosis. ELISA quantification of secreted SPP1(OPN) levels in murine YTN16 cell lines, peritoneal tumor cells and macrophages cultured at equivalent densities. N=3 per group \*\*\**p<0.001*, \*\*\*\**p< 0.0001*, by one-way ANOVA was used to compare the differences between groups.

SPP1 has been shown to be expressed by tumoral and stromal compartments. It has been described as a marker of macrophage state, highly expressed by M2-like macrophages^17–20^. Given the high levels of SPP1 secretion by YTN16 cells, we posited that tumor cells are a main source of secreted SPP1 in gastric peritoneal carcinomatosis. Single-cell RNA sequencing of human malignant ascites revealed that multiple cellular populations expressed *SPP1* transcripts (Supplemental Figure 1C); however, these data do not distinguish which cells actively secrete the protein. To quantitatively determine the cellular sources of SPP1, we performed ex-vivo profiling of sorted CD45^-^Epcam^+^ tumor cells and macrophages isolated from peritoneal metastatic nodules in tumor bearing mice. Quantitative protein analysis by ELISA demonstrated that tumor cells were the predominant source of secreted SPP1, producing higher levels than peritumoral macrophages (Figure 2I).

### SPP1 promotes gastric peritoneal carcinomatosis

The functional role of SPP1 in gastric cancer peritoneal dissemination has not been previously described. Analysis of patient datasets revealed that elevated *SPP1* expression was associated with reduced overall survival and progression free survival in gastric cancer, a trend that persisted among patients with Stage IV metastatic disease (Figure 3A-B). Utilizing our murine model of gastric peritoneal carcinomatosis, intraperitoneally administered pharmacologic inhibition of SPP1 markedly restricted tumor progression, resulting 40% reduction in tumor weight and 50% decrease in peritoneal nodules (Figure 3C). Consistent with these findings, inhibition of SPP1 profoundly altered the peritumoral inflammatory milieu. Analysis of the whole tumor secretome by Luminex demonstrated that levels of the chemoattractant CCL2, which recruits macrophages and tolerogenic IL-10 were diminished compared to controls. In contrast, inflammatory mediators associated with enhanced T-cell responses, including IL-12 and CCL4, were increased (Figure 3D). These findings establish SPP1 as a critical driver of peritoneal dissemination and immunosuppression in gastric peritoneal carcinomatosis.

**Figure 3:**
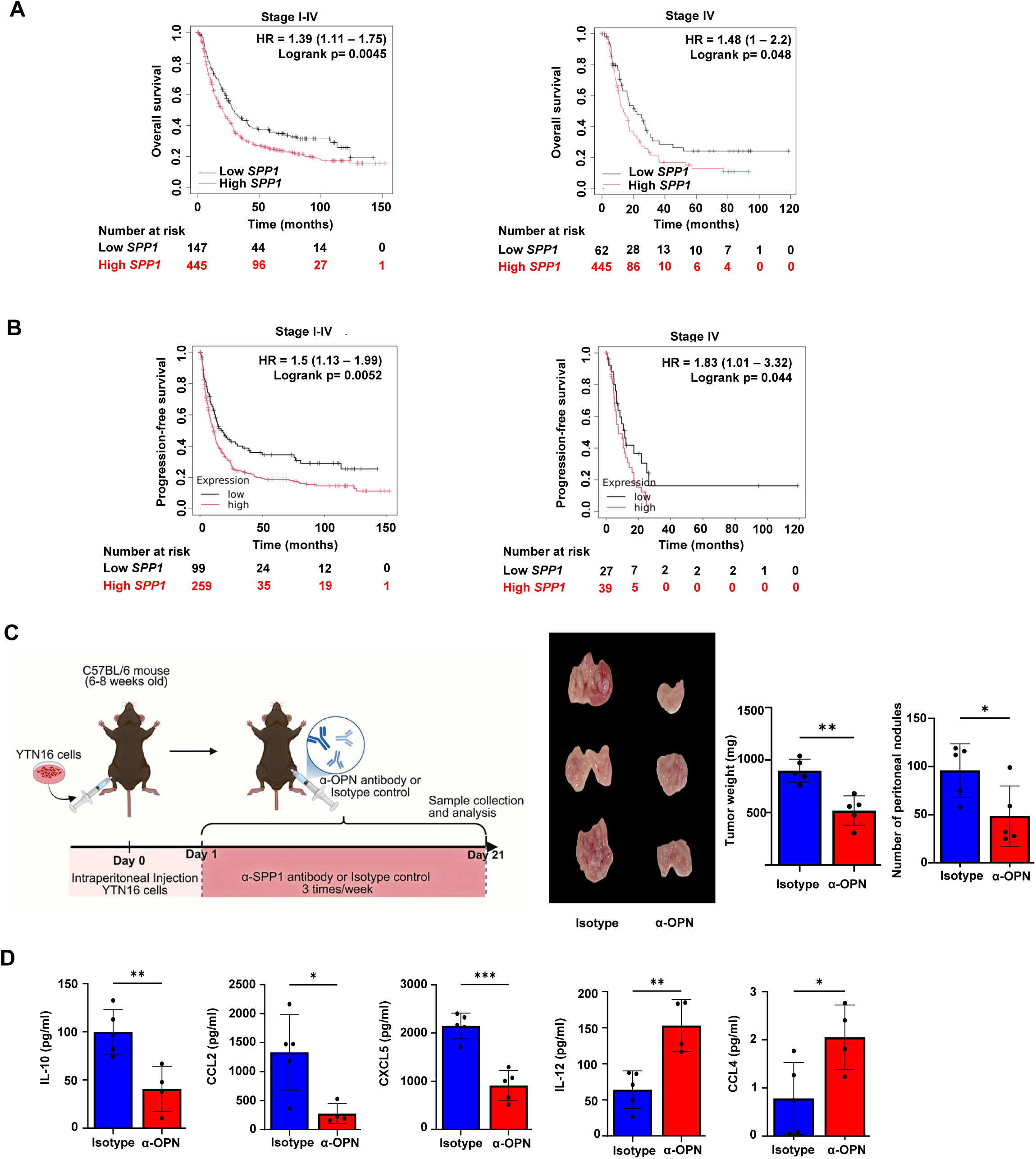
Intracavitary SPP1 blockade inhibits gastric cancer peritoneal dissemination. (A) Kaplan-Meier curves showing overall survival of gastric cancer patients stratified by SPP1 mRNA expression (B) Kaplan-Meier curves showing progression free survival of gastric cancer patients stratified by SPP1 mRNA expression. (C) Schematic of treatment paradigm of SPP1(OPN) blocking antibody in gastric peritoneal carcinomatosis in mice. Intraperitoneal (IP) injection of SPP1 blocking antibody in tumor bearing mice. Omental tumor weight and total number of tumor nodules depicted. N=5 per group (D) Quantification of IL-10, CCL2, CXCL5, IL-12 and CCL4 levels from the tumor secretome by Luminex assay. N=5 per group \**p< 0.05*, \*\**p< 0.01*, \*\*\**p < 0.001*, by unpaired t test was used to compare the differences between groups. HR=Hazard Ratio

### Paracrine effects of tumor-secreted SPP1 promote macrophage recruitment

Because we observed that SPP1 inhibition decreased the macrophage chemoattractant CCL2 (Figure 3D), we sought to determine whether SPP1 directly regulates macrophage recruitment. Analysis of TIMER datasets demonstrated a positive association between macrophage infiltration and *SPP1* transcript levels in gastric cancer (Figure 4A). Consistent with this observation, gastric cancer patients harboring *SPP1* mutations exhibited reduced macrophage infiltration relative to patients lacking *SPP1* alterations (Figure 4B). Functionally, recombinant SPP1 promoted macrophage migration in vitro in a dose-dependent manner (Figure 4C). In-vivo, pharmacological blockade of SPP1 (OPN) significantly reduced peritumoral macrophage infiltration in the mouse model of gastric peritoneal carcinomatosis (Figure 4D). Together, these findings, coupled with the observation that tumor cells are the predominant source of SPP1 (Figure 2I) demonstrate that tumor derived SPP1 selectively promotes pro-tumorigenic macrophage recruitment within the peritoneal tumor microenvironment, thus enhancing tumor dissemination.

**Figure 4:**
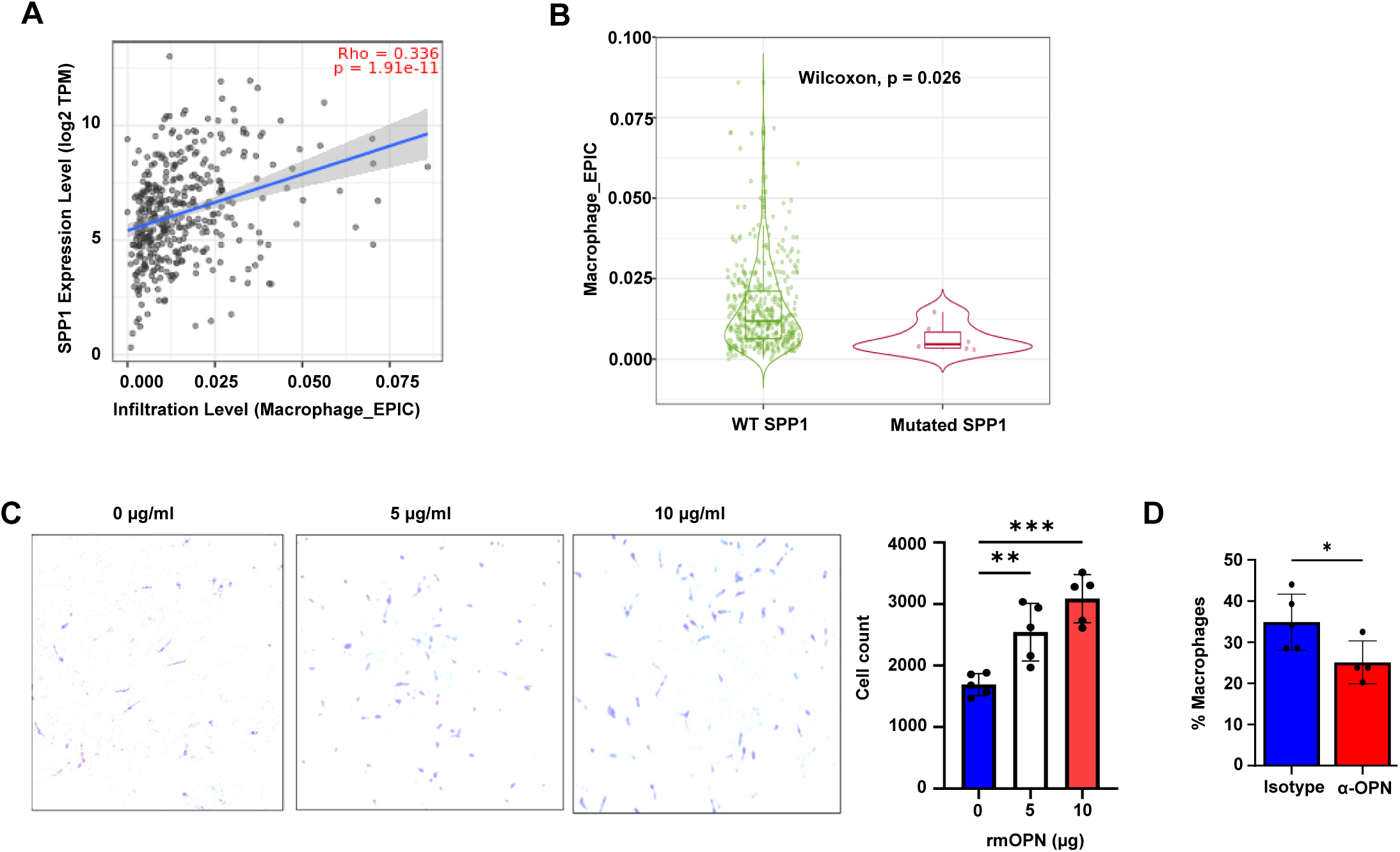
SPP1 drives macrophage recruitment into the peritoneal tumor microenvironment. (A) Correlation between SPP1 expression and macrophage infiltration in gastric cancer patients analyzed by TIMER (B) Comparison of macrophage infiltration between SPP1-mutant and SPP1 wild-type gastric cancer patients analyzed by TIMER. (C) In vitro BMDM migration assay following treatment with recombinant SPP1 (rmOPN) compared to BSA carrier. N=5 per group (D) Flow cytometric quantification of tumor-infiltrating macrophages in peritoneal metastasis following *in vivo* SPP1 blockade compared with isotype control treatment. *\*p< 0.05*, \*\**p<0.01*,\*\*\**p< 0.001*, by unpaired t-test for experiments with two groups and one-way ANOVA for experiments with three groups.

### Macrophage depletion diminishes peritoneal dissemination

To test the relevance of macrophage infiltration in the metastatic microenvironment, Immunohistochemical analysis of human gastric carcinomatosis specimens revealed enrichment of CD68⁺ macrophages compared with normal peritoneal omental tissue (Supplemental Figure 2A). Consistent with this observation, single-cell RNA sequencing of human malignant ascites demonstrated substantial macrophage infiltration (Supplemental Figure 2B). To model these findings in vivo, we utilized the aggressive murine model of peritoneal carcinomatosis by intraperitoneal injection of syngeneic YTN16 gastric tumor cells (Figure 2E). Flow cytometric analysis revealed marked enrichment of Gr1⁻CD11b⁺ F4/80⁺ macrophages in peritoneal metastases and malignant ascites compared with control spleen (Supplemental Figure 2C).

Based on these observations, we hypothesized that macrophages promote tumor progression in gastric peritoneal carcinomatosis. Consistent with this hypothesis, macrophage depletion using clodronate liposomes phenocopied the anti-tumor effects of SPP1 blockade, significantly reducing peritoneal tumor burden, metastatic nodule formation, and manifestations of malignant intestinal obstruction (Figure 5A-E). Accordingly, analysis of the whole tumor secretome by Luminex demonstrated that levels of IL-10 and associated cytokines IL-6, IL-22 and IL-17A were inhibited by macrophage depletion (Figure 5F)

**Figure 5:**
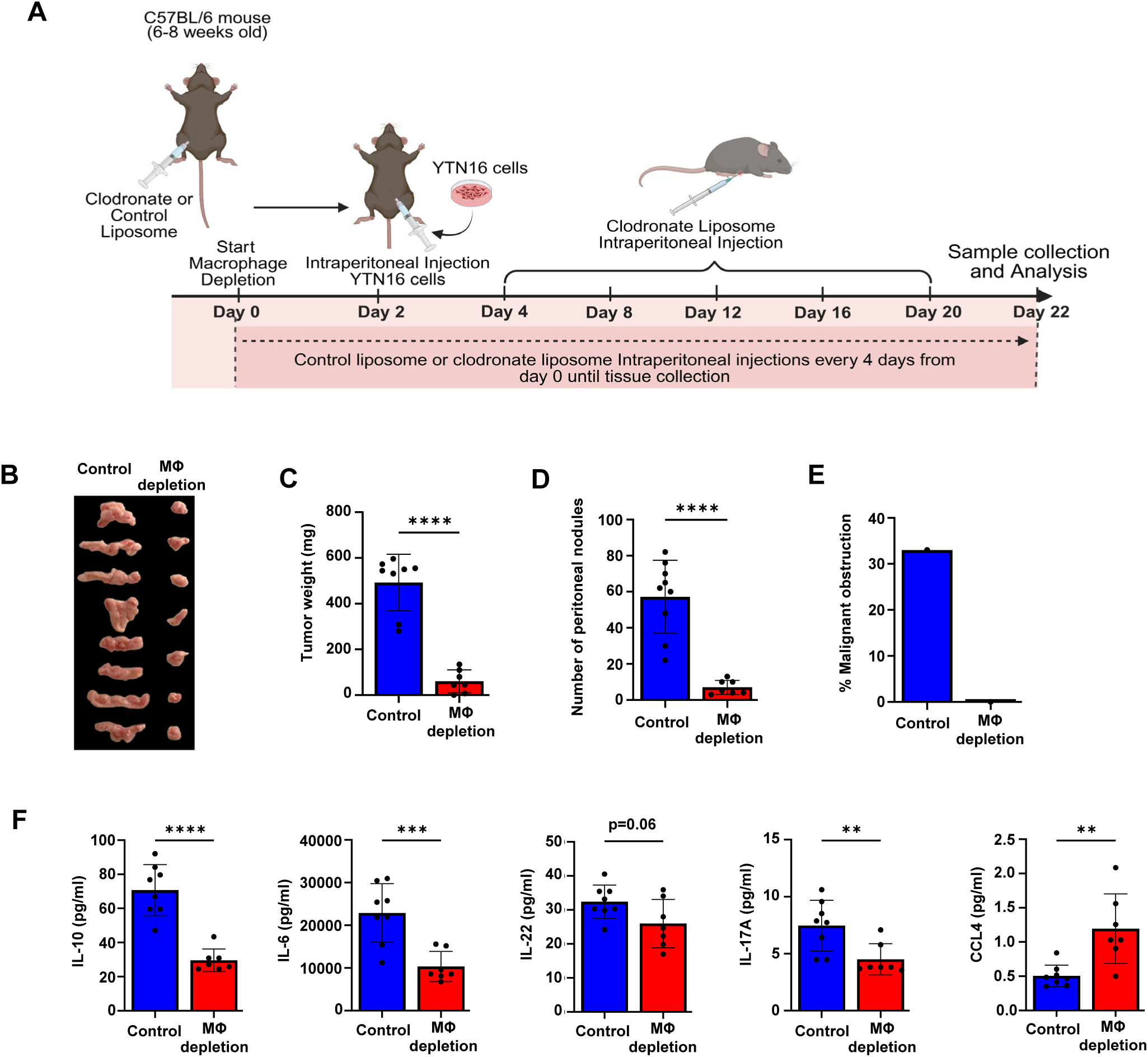
Intracavitary macrophage depletion inhibits gastric cancer peritoneal dissemination and alters inflammatory milieu. (A) Schematic of in-vivo macrophage (MՓ) depletion using clodronate liposomes (B-E) Omental tumor weights, number of total peritoneal nodules, and symptomatic malignant obstruction with macrophage depletion N=8 per group (F) Quantification of IL-10, IL-6, IL-22, IL-17A and CCL4 levels from the tumor secretome by Luminex assay. N=8 per group \*\**p< 0.01*; \*\*\**p< 0.001*, \*\*\*\**p<0.0001*, by unpaired t test was used to compare the differences between groups.

### Tumor derived SPP1 induces tolerogenic IL-10 macrophage secretion

Given that both SPP1 inhibition and macrophage depletion in tumor-bearing mice decreased IL-10 levels (Figure 3D, 5F), we next investigated whether tumor derived SPP1 promotes immunoregulatory macrophage programming through induction of IL-10 secretion. Consistent with this idea, analysis of TIMER datasets revealed a positive correlation between *SPP1* and *IL10* transcript expression in gastric cancer (Figure 6A). To infer tumor-macrophage communication networks, NicheNet analysis was performed on patient-derived malignant ascites. This in-silico approach predicts how ligands produced by one cell population influence gene programs in another. This analysis identified *SPP1* tumor cell ligand predicted to interact with macrophage surface receptors and induce downstream regulation of *IL10* expression (Figure 6B-C). Notably, macrophages emerged as the predominant source of *IL10* among immune cell populations within the ascites (Figure 6D). Mechanistically, recombinant SPP1 directly stimulated IL-10 secretion by murine macrophages in-vitro (Figure 6E). Together, these findings indicate that tumor-derived SPP1 exerts immunomodulatory effects by both promoting macrophage recruitment and driving their polarization toward a tumor-permissive phenotype within the peritoneal tumor microenvironment.

**Figure 6:**
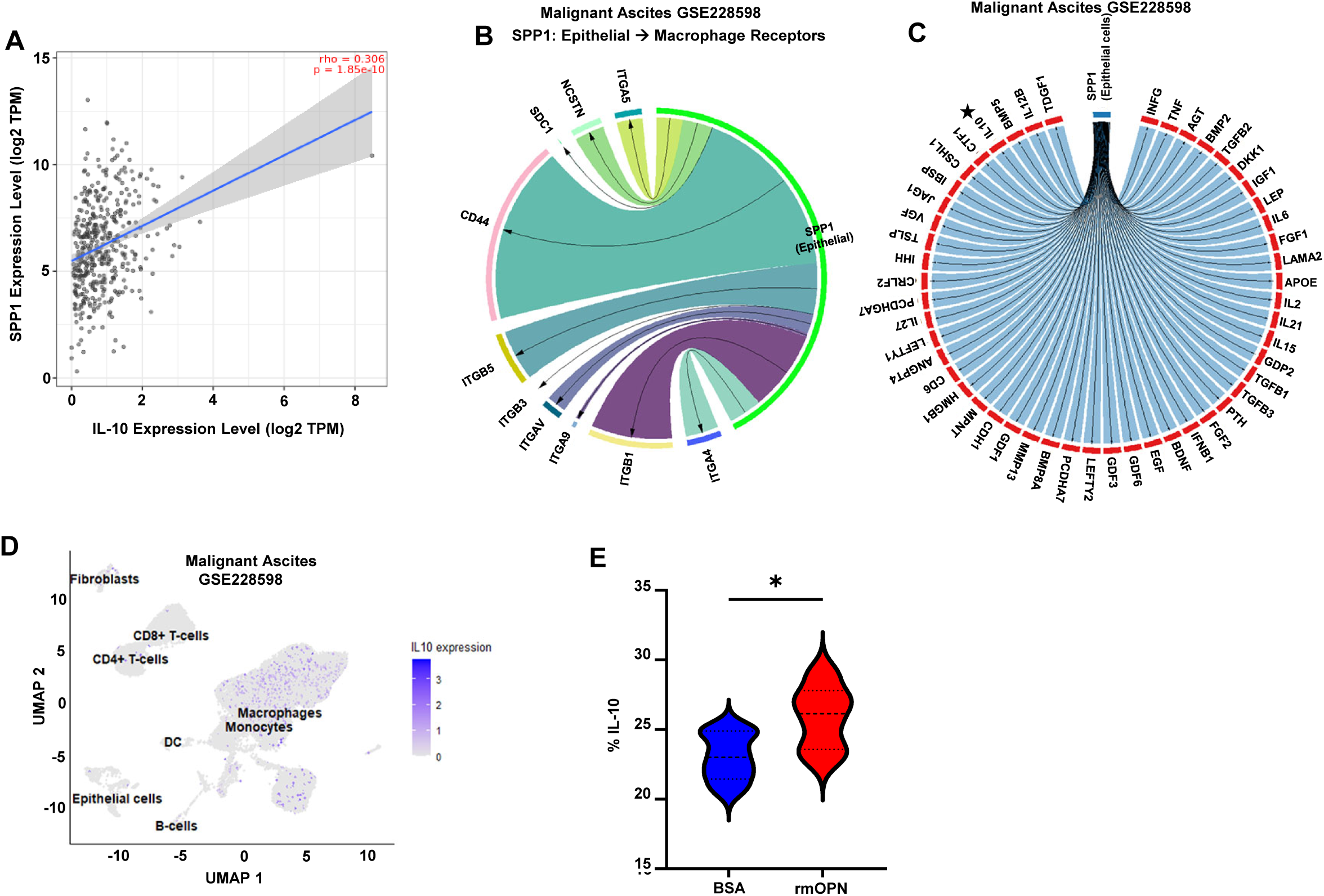
Tumor derived SPP1 induces macrophage IL-10 secretion. (A) Correlation between IL10 expression and macrophage infiltration in gastric cancer patients analyzed by TIMER (B)NicheNet analysis from single cell sequencing of human gastric cancer malignant ascites predicting epithelial-derived SPP1 ligand interactions with cognate receptors expressed on macrophages. (C) NicheNet analysis from single cell sequencing of human gastric cancer malignant ascites predicting downstream macrophage transcriptional programs associated with epithelial-derived SPP1 signaling. (D) Single cell sequencing showing IL10 expression in human gastric cancer malignant ascites across cell types. (E) In-vitro flow cytometric analysis of IL-10 expression BMDM (MՓ) following treatment with recombinant SPP1(rmOPN) compared to BSA carrier control. BMDM activated with IL-4 and IL-13 in both groups. N=5 per group. \**p<0.05*, by unpaired t test was used to compare the differences between groups.

## Discussion

In this study, we identify a novel tumor-macrophage signaling axis that promotes metastatic progression in gastric peritoneal carcinomatosis. Leveraging a multi-omic approach integrating human specimens and murine models of peritoneal metastasis, we demonstrate that macrophages accumulate within the peritoneal metastatic niche and adopt immunosuppressive phenotypes that promote tumor growth. We further show that gastric tumor-derived SPP1 drives macrophage recruitment and induces IL-10-dependent immunosuppressive programming. Functionally, SPP1 promotes peritoneal tumor dissemination. Together, these findings suggest that therapeutic disruption of this tumor-macrophage signaling axis may restore anti-tumor immunity and limit disease progression.

Tumor-secreted factors are increasingly recognized as a fundamental hallmark of cancer that enables tumors to remodel their microenvironment, evade immune surveillance, and support metastatic progression. Among these factors, SPP1 has increasingly been recognized as an important regulator of tumor progression across multiple malignancies. In pancreatic cancer, SPP1 has been shown to regulate epithelial-mesenchymal plasticity and tumor cell fate through paracrine signaling networks^21^ while studies in breast^32^, colon^33^ and esophageal^34^ cancer have demonstrated that tumor-derived SPP1 promotes macrophage reprogramming. Despite being identified as an independent predictor of peritoneal recurrence in gastric cancer^22^, the functional role of tumor-derived SPP1 within the gastric peritoneal metastatic microenvironment has remained largely unexplored. This represents a critical knowledge gap given the central role of the peritoneum as a site of gastric cancer dissemination and the established association between pro-tumorigenic macrophages and disease progression^13^. Our findings demonstrate that tumor-derived SPP1 promotes macrophage recruitment and immunoregulatory reprogramming, supporting the development of a pro-metastatic niche and further highlighting SPP1 as a potential therapeutic target in gastric peritoneal carcinomatosis.

Transcriptomic analyses across cancers including colorectal, head and neck squamous cell carcinoma, and non-small cell lung cancer have similarly identified SPP1-expressing macrophages as a distinct immunosuppressive subset within the tumor microenvironment^17–20^. In contrast to studies emphasizing macrophage-derived SPP1, our findings identify tumor cells as a major source of SPP1 in gastric peritoneal carcinomatosis and demonstrate that tumor-secreted SPP1 actively promotes macrophage recruitment and IL-10 dependent immunosuppressive programming. These results suggest that tumor-derived SPP1 functions as an upstream regulator of macrophage-mediated immune suppression, establishing an immunosuppressive metastatic niche that supports peritoneal tumor dissemination. Our findings broaden the perspective of SPP1 from a marker of macrophage states to a tumor-derived signal that actively orchestrates macrophage recruitment and reprogramming, which in turn supports immunosuppressive niche formation during metastatic progression.

We have identified the SPP1-macrophage axis as a potential therapeutic strategy in gastric peritoneal carcinomatosis. Intraperitoneal delivery is particularly compelling, given the limited efficiency of systemic therapies in penetrating the blood-peritoneal barrier^35,36^. In addition to targeting SPP1, we demonstrate that local intraperitoneal modulation of macrophages is sufficient to restrict peritoneal dissemination, highlighting the central role of macrophages in driving metastatic progression. While macrophage-targeted therapies raise concerns regarding systemic myelosuppression^37^, our results show that iterative localized intraperitoneal delivery is both well tolerated and effective in preclinical models. Future studies will focus on combinatorial immunotherapies and chemotherapies to eradicate peritoneal carcinomatosis.

While our study establishes a tumor-macrophage signaling axis that drives metastatic progression, several limitations warrant further investigation. We do not comprehensively describe the full spectrum of macrophage heterogeneity within the peritoneal tumor microenvironment. Future studies will build on these findings by delineating functionally distinct macrophage subsets that drive peritoneal dissemination and by defining the upstream signaling pathways that mediate SPP1-induced macrophage programming.

Our findings suggest that gastric tumor-derived SPP1 represents an upstream driver of macrophage-mediated immune suppression. These findings highlight the central role of tumor derived factors in shaping the innate immunity and extend the functional relevance of SPP1 beyond a macrophage associated marker to an upstream regulator of macrophage programming. Our results provide a strong rationale for targeting the SPP1-macropage axis as a novel therapeutic target in this otherwise refractory disease. More broadly, this work underscores the pivotal importance of tumor-immune crosstalk in metastatic progression and suggests that harnessing the immunomodulatory properties of tumor cells may be a generalizable therapeutic approach across various cancers.

## Methods

### Sex as a biological variable

Male and female C57BL/6J mice between six- to eight-month-old were used in all experiments. Similar findings are reported for both sexes.

### Statistics

Data is presented as mean +/- standard error. Statistical significance was determined by the Student’s unpaired t-test for comparison of two groups and analysis of variance (ANOVA) for multiple comparisons using GraphPad Prism 7 (GraphPad Software, La Jolla, CA). p-values <0.05 were considered significant. Data represent mean ± SEM.

### Study approval

All animal experiments were performed in accordance with the University of Kentucky Institutional Animal Care and Use Committee (IACUC) guidelines. De-identified human tissue specimens were obtained through the University of Kentucky Biospecimen Procurement and Translational Pathology Shared Resource under a protocol approved by the University of Kentucky Institutional Review Board (IRB protocol #48495). Publicly available, de-identified datasets were obtained from the Gene Expression Omnibus (GEO), The University of Alabama at Birmingham Cancer data analysis Portal (UALCAN), TIMER, The Cancer Genome Atlas (TCGA), and the Genotype-Tissue Expression (GTEx) project and did not require additional IRB approval.

### Data availability

The single-cell RNA sequencing dataset analyzed in this study is publicly available through the Gene Expression Omnibus (GEO) under accession number GSE228598. Publicly available transcriptomic and clinical datasets were obtained from TCGA-STAD, GCR-STAD, ACRG/GSE66229, GSE162214, GTEx, TIMER2.0, UALCAN and KM Plotter all described below. The murine bulk RNA sequencing data generated in this study will be deposited in the Gene Expression Omnibus (GEO), and the accession number will be provided upon acceptance. All other data supporting the findings of this study are available from the corresponding author upon reasonable request.

### Human data source and transcriptomic processing

Transcriptomic data for gastric adenocarcinoma were obtained from publicly available datasets. *SPP1* mRNA expression was compared between normal and tumor tissues across datasets. TCGA-STAD and ACRG^23,24^ (GSE66229, GSE162214) were obtained through the UCSC Xena platform https://xena.ucsc.edu. Normal stomach tissue expression data were obtained from the GTEx portal https://gtexportal.org. UALCAN^25^ https://ualcan.path.uab.edu/ and the Gastric Cancer Registry (GCR) https://gcregistry-explorer.stanford.edu/ were also utilized. Immune infiltration and genomic correlation analyses were performed using TIMER3.0^26^ https://compbio.cn/timer3/, and Kaplan-Meier analyses for overall and progression-free survival were generated using the Kaplan-Meier Plotter^27^ platform https://kmplot.com/analysis.

### Single-cell RNA sequencing

Single-cell RNA sequencing data from malignant ascites of patients with gastric cancer were obtained from the Gene Expression Omnibus (GEO; GSE228598)^28^, and processed gene expression matrices from 15 samples were integrated into a single Seurat object using Seurat (v5.1.0) in R (v4.5.2). Dimensionality reduction was performed using principal component analysis, and clusters were visualized by Uniform Manifold Approximation and Projection (UMAP); cluster-specific marker genes were identified using *FindAllMarkers* with a log2 fold-change threshold of 0.25 and a minimum expression threshold of 10% of cells per cluster. Cell type annotation was performed using SingleR, and epithelial tumor cell-macrophage ligand-receptor interactions were inferred using NicheNet.

### Animals and In Vivo Procedures

C57BL/6J mice (Jackson Laboratory) were housed in a pathogen-free environment and fed standard mouse chow at libitum. All animal experiments were performed in accordance with the University of Kentucky Institutional Animal Care and Use Committee (IACUC) guidelines. Sex-and age-matched C57BL/6J mice between six- to eight-month-old were used in all experiments.

Gastric peritoneal carcinomatosis was established by intraperitoneal injection of 1 x10^6^ YTN16 cells in PBS, and mice were sacrificed 21 days after implantation or upon reaching humane endpoints. For in vivo antibody studies, mice were randomized one day after tumor implantation and treated intraperitoneally every 72 hours with 200 μg of neutralizing anti-SPP1 antibody (BioXcell, 103D6) or isotype control antibodies (BioXcell, DV5-1). For macrophage depletion, clodronate or control liposomes (F70101C-NC-10, FormuMax) were administered intraperitoneally beginning 2 days before tumor implantation and continued every 4 days throughout the experiment.

### Cell lines

Syngeneic murine gastric YTN16 cells^29^ were maintained in complete medium consisting of Dulbecco’s Modified Eagles Medium (DMEM) supplemented with 10% heat-inactivated FBS and penicillin/streptomycin. RAW264.7 (TIB-71, ATCC) were maintained in complete DMEM.

### Conditioned media (CM) and Recombinant SPP1 macrophage culture

3 x10^6^ YTN16 cells were cultured until 70% confluent. CM was collected in a 50-ml conical tube and centrifuged. Supernatant was carefully transferred to a new tube and used for assays and diluted 1:1 with control media for macrophage culture experiments.

Bone marrow-derived macrophages (BMDM) were isolated from femurs of C57BL/6J mice as previously described^30,31^. After isolation, BMDMs were incubated in 50 ng/ml M-CSF (Miltenyi Biotec). On day 7, BMDM were seeded in LPS (100 ng/ml, Sigma) or IL-4 (20 ng/ml, Miltenyi Biotec) and IL-13 (20 ng/ml, Miltenyi Biotec) for 24-48 hours. In select experiments, recombinant mouse OPN (SPP1) (R&D System) or bovine serum albumin (BSA) was added to BMDMs activated with IL-4 (20 ng/ml, Miltenyi Biotec) and IL-13 (20 ng/ml, Miltenyi Biotec) for 24-48 hours. RAW264.7 macrophages were similarly treated with conditioned or control media.

### Cytokine Array and Multiplex assays

Conditioned media from YTN16 was collected for Mouse Cytokine Array C1000 (RayBiotech, Catalog# AAM-CYT-1000-4) following manufacturer’s instructions. For quantitative multiplex assays, tumor pieces of 8 mg (±0.3 mg) were incubated in DMEM for 18 hours. Tumor culture supernatants were analyzed by Procarta multiplex bead array. The assay was run according to manufacturer’s instructions on a Luminex LX200 instrument.

### Transwell migration assay

BMDM were seeded onto 8 µm pore insert placed in a 24-well companion plate. Recombinant mouse OPN (OPN) or BSA carrier as control were added in lower chamber and incubated for 6hours. Slides were analyzed under Nikon microscope and images analyzed with ImageJ software.

### Western blot

Tumor tissues and cell lysates were prepared in RIPA buffer supplemented with freshly added protease and phosphatase inhibitors. Tumor tissues were homogenized in RIPA buffer (10 μL/mg tissue) with zirconium ceramic oxide beads (Fisher brand). Protein lysates (10–40 μg) were separated by SDS-PAGE on 4–20% precast gels (Bio-Rad), transferred to PVDF membranes, and incubated with antibodies against SPP1 and GAPDH. Protein bands were detected using conjugated secondary antibodies and enhanced chemiluminescence, visualized on a ChemiDoc Imaging System, and quantified using Image Lab software (Bio-Rad).

### RNA isolation and RTqPCR

Total RNA was extracted using the miRNeasy Tissue/Cells Advanced Mini Kit or TRIzol reagent according to the manufacturers’ instructions. RNA concentration was quantified using a NanoPhotometer. Equal amounts of RNA were reverse transcribed into cDNA using the High-Capacity cDNA Reverse Transcription Kit. Quantitative real-time PCR was performed in duplicate using SYBR Green chemistry on a QuantStudio 6 Flex Real-Time PCR System, and relative gene expression was calculated using the comparative threshold cycle (2^-ΔΔCt) method normalized to Gapdh. Primer sequences *Arg1* gene specific primer-Forward: 5’ GTGAAGAACCCACGGTCTGT 3’ and *Arg1* Reverse: 5’ CTGGTTGTCAGGGGAGTGTT 3’; *Tnfα* gene specific primer -Forward: 5’ TGCCTATGTCTCAGCCTCTTC 3’ and *Tnfα*-Reverse: 5’ GGTCTGGGCCATAGAACTGA 3’; *IL1β* gene specific primer -Forward: 5’ GCAACTGTTCCTGAACTCAACT 3’ and *IL1β*-Reverse: 5’ ATCTTTTGGGGTCCGTCAACT 3’; *Nos2* gene specific primer -Forward: 5’ GCAGCTGGGCTGTACAAA 3’ and *Nos2*-Reverse: 5’ AGCGTTTCGGGATCTGAAT 3’.

### Flow cytometry and Cell sorting

Malignant ascites and peritoneal tumors were harvested from tumor-bearing mice, and single-cell suspensions were prepared as previously described with minor modifications^9,10^. Peritoneal tumors were mechanically dissociated and enzymatically digested in HBSS containing 2% heat-inactivated FBS, collagenase IV, and DNase I at 37°C for 20 minutes, followed by filtration through a 40μm mesh and centrifugation. Malignant ascites was obtained by injection of PBS intraperitoneally and needle aspirated. Splenocytes were isolated by mechanical disruption, and all samples underwent red blood cell lysis before viability staining and Fc receptor blockade with anti-CD16/CD32 antibody. After live/dead discrimination with Zombie Yellow, cells were stained with fluorescently conjugated antibodies for surface markers CD45, CD11b, Gr1, F4/80, CD86 and MHCII. For intracellular staining, cells were fixed and permeabilized and incubated with Arg-1, TNF-α and IL-10. Flow cytometry data were acquired using a FACSymphony A3 cell analyzer (BD Biosciences) and analyzed with FlowJo software.

For cell sorting, single-cell suspensions from tumors were stained with fluorescently conjugated antibodies. Cells were resuspended in sorting buffer and viable cells were identified by exclusion of Propidium Iodide. Cell sorting was performed using a FACSymphony S6 Cell Sorter (BD Biosciences), and sorted cells were cultured for 48 hours. Supernatants were collected, and SPP1 levels were quantified by ELISA (Mouse/Rat Osteopontin/OPN ELISA Kit, catalog# KE10118) according to the manufacturer’s instructions.

### Bulk RNA sequencing

BMDMs, malignant ascites, and peritoneal tumor tissues were collected processed for RNA isolation as described above. Total RNA was used for library preparation with the Illumina Stranded mRNA Prep kit according to the manufacturer’s instructions. Raw RNA-sequencing reads were aligned to the *Mus musculus* reference genome (GRCm38/mm10) using STAR, and sequencing quality metrics, transcript coverage, and strand specificity were assessed using RSeQC. Gene-level read counts were generated using HTSeq-count, normalized with DESeq2, and differentially expressed genes were identified based on fold change and adjusted *P* value thresholds. Genes with log2Fold Change ≥2 and adjusted *p value <0.05* were considered significant. Functional enrichment analyses, including Gene Ontology ORA was performed to characterize significantly upregulated biological pathways (adjusted *p value <0.05*). Predicted pathway activity scoring based on z-scored gene expression was also performed.

### Immunohistochemistry

Human gastric peritoneal metastasis and normal omental tissue slides obtained from formalin fixed paraffin embedded blocks were incubated with anti-CD68 (Clone KP-1, Roche 05278252001) followed by OmniMap anti-Mouse-HRP (Roche 05269652001.

## Author contributions

Conceptualization- MBH

Methodology- MBH, LT, NH, MC, EB, EU

Formal Analysis- MBH, LT, NH, EB

Resources- SN, EB, MBE, YJ, JK, JL, MBH

Visualization-LT, NH, MBH

Writing- LT, NH, MBH

## Acknowledgements

We thank the Biospecimen Procurement & Translational Pathology Shared Resource Facility of the University of Kentucky Markey Cancer Center (supported by National Cancer Institute grant P30 CA177558).

This work was supported by the UK Flow Cytometry & Immune Monitoring core facility. (RRIDSCR_026358)

Murine YTN16 cell line was a gift of Dr. Ellen Beswick and Dr. Sachiyo Nomura

