## Supplemental Figure 1 for "Tumor-Derived SPP1 Drives Immunosuppressive Macrophage Reprogramming in Gastric Peritoneal Carcinomatosis"

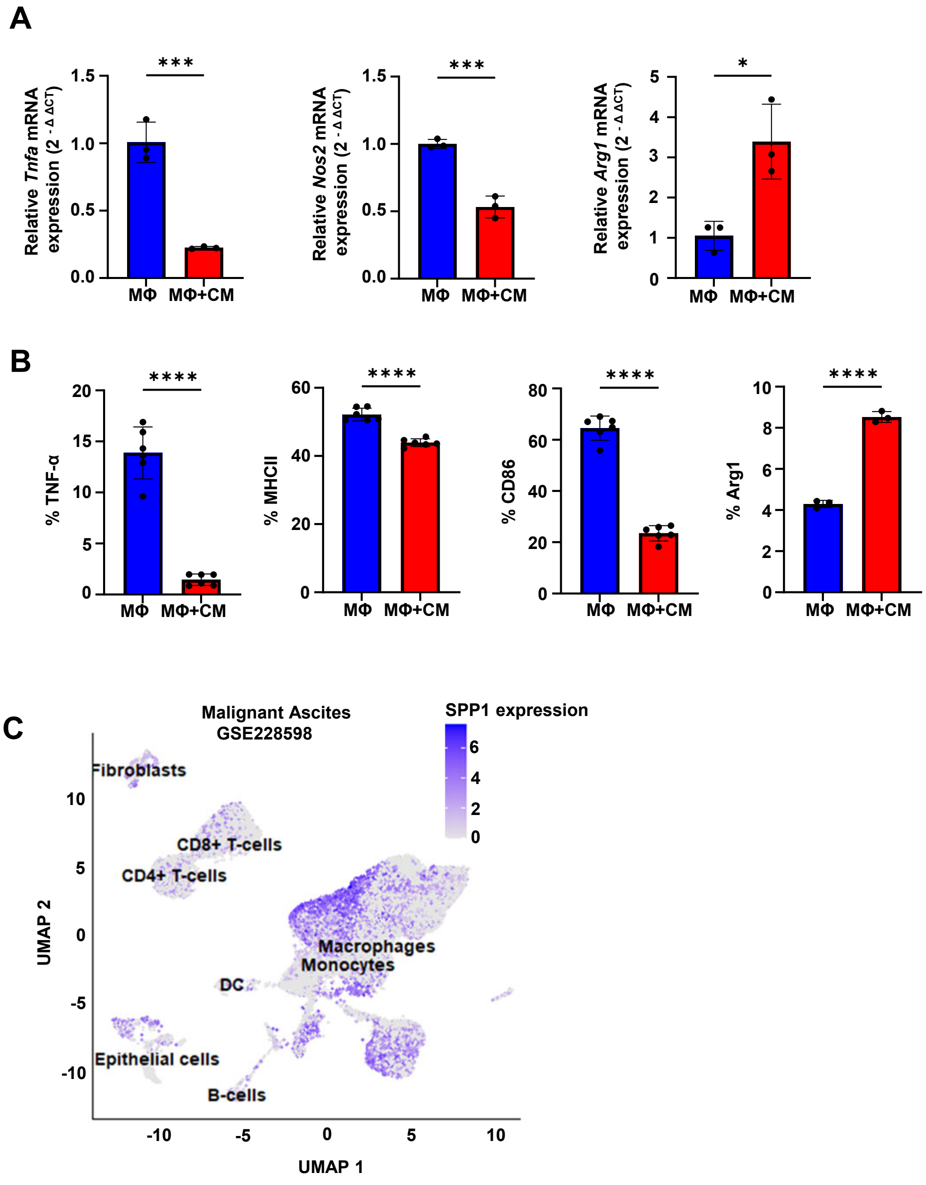


**Supplemental Figure 1**: **Gastric cancer secreted factors promote an anti-inflammatory macrophage phenotype**. (A) RT-qPCR of *Arg1*, *Tnfa*, *Nos2* in RAW 267.4 macrophages (MΦ) treated with conditioned media (CM) compared with untreated macrophages (MΦ). N=3 per group (B) Flow cytometry showing protein levels of TNF-α, MHCII, CD86 and Arg-1 in RAW 267.4 macrophages incubated in CM compared with untreated MΦ. RAW 267.4 macrophages were activated with LPS in both conditioned media and untreated groups. N=5 per group. (C) Single cell sequencing showing IL10 expression in human gastric cancer malignant ascites across cell types. *$p<0.05$, **$p< 0.01$; ***$p< 0.001$, ****$p<0.0001$, by unpaired t test was used to compare the differences between groups.
