## Supplemental Figure 2 for "Tumor-Derived SPP1 Drives Immunosuppressive Macrophage Reprogramming in Gastric Peritoneal Carcinomatosis"

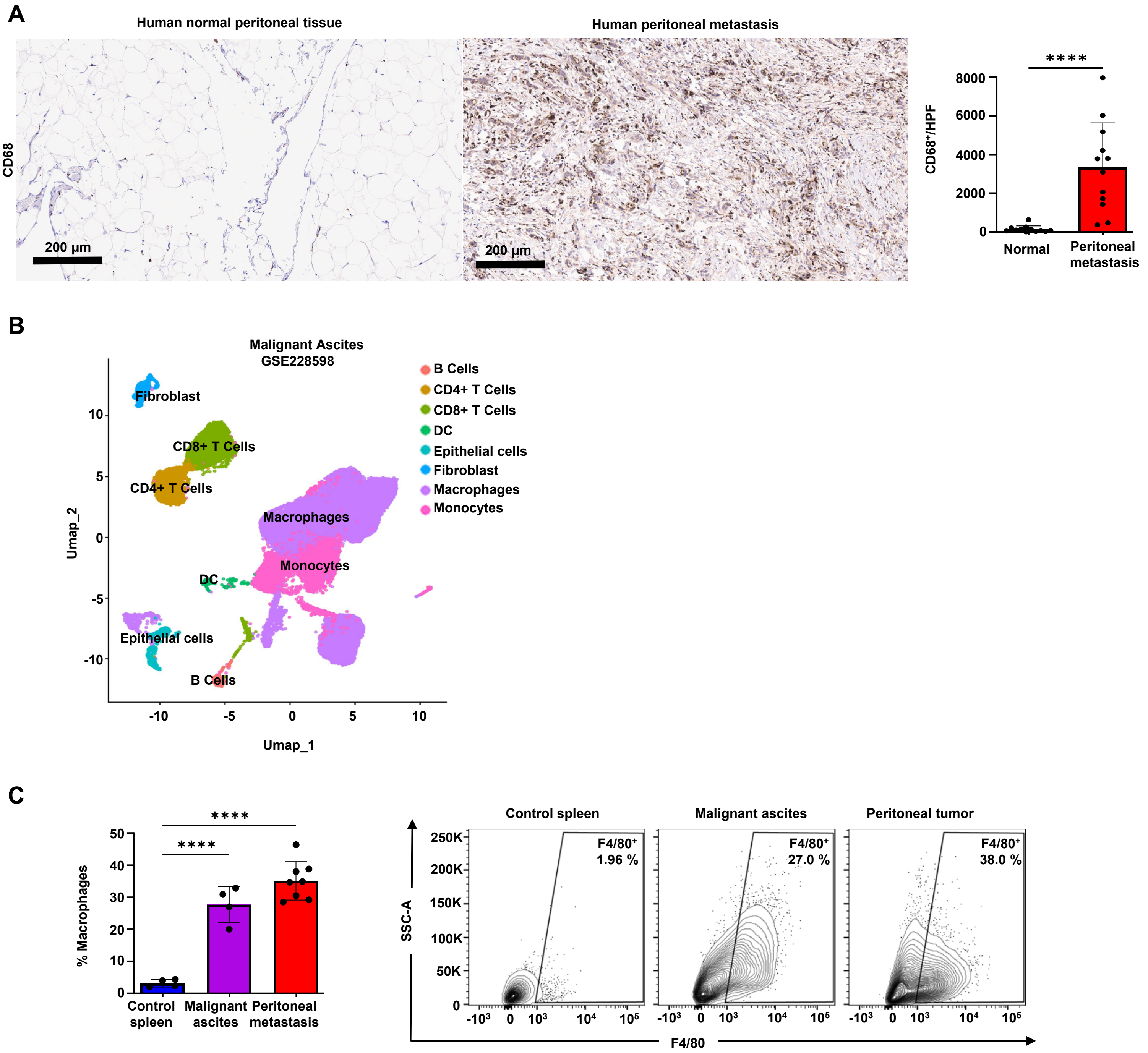


**Supplemental Figure 2: Macrophages are an abundant cell population in human and mice gastric peritoneal carcinomatosis.** (A) Immunohistochemistry of $CD68^{+}$ macrophages of human gastric peritoneal metastasis and normal omental tissue. N=12 per group. (B) Single cell sequencing of cell populations in human gastric malignant ascites. (C) Flow cytometric analysis of $CD11b^{+}Gr1^{-}F4/80^{+}$ macrophages in matched murine malignant ascites and peritoneal metastasis compared with spleen in tumor bearing mice. ****$p<0.0001$, by unpaired t-test for experiments with two groups and one-way ANOVA for experiments with three groups.
